# Antibiotic resistance genes in sewages from hospitals and the urban setting in a Peruvian city in the highlands

**DOI:** 10.1101/2022.12.13.520267

**Authors:** Julio A. Poterico, Luis Jaramillo-Valverde, Nelis Pablo-Ramirez, Vicky Roa-Linares, Catalina Martinez-Jaramillo, Sandra Alvites-Arrieta, Milward Ubillus, Diana Palma-Lozano, Rony Castrejon-Cabanillas, Samuel Davison, A. Gomez, Heinner Guio

## Abstract

**Background:** The establishment of metagenomics seems a suitable approach to assess the abundance and diversity of antibiotic resistance genes (ARG)

**Methods:** Metagenomics study in a Peruvian city from the highlands, where samples were derived from sewage waters from two hospitals and the urban setting. DNA extraction was performed in 250 mL and then 16S rRNA gene amplification and shotgun sequencing were carried out. The bioinformatics pipeline was performed following recommendations for metagenomics analysis. Alpha diversity was evaluated with the Shannon and Simpson’s indices; whereas beta diversity was evaluated through the Bray-Curtis index, and using the principal coordinate analysis (PCoA) to explore and visualize the differences.

**Results:** We found a high abundance of bacteria related to resistance to beta-lactams, macrolides, aminoglycosides, fluoroquinolones, and tetracyclines. The urban sample did not differ significantly from the wastewater ARG presence from the hospitals in Huanuco.

**Conclusion:** Metagenomics analysis through sewage strategies seems to help to monitor the AMR to establish local public health policies, especially in cities or countries with limited resources to establish large projects conceiving the One Health approach.

## INTRODUCTION

The current policy framework of the World Health Organization (WHO) relies on the One Health concept. This notion involves not only the human and animal contribution to global health but the role of the environment. The interaction between human-to-human, animal-to-animal, human-to-animal, and the environment with the aforementioned, fosters the development of antimicrobial resistance (AMR) (1).

AMR represents a global health threat due to several drivers contributing to superbug emergence. For instance, Murray et al. reported that 4.95 million people had AMR-related health problems in 2019(2), with six pathogens directly associated with AMR deaths. Furthermore, the daunting AMR projections propose that by 2050, one person would die every three seconds, in other words, ten million people would die due to AMR (3).

The WHO established 2015 the Global Antimicrobial Resistance and Use Surveillance System (GLASS), a global strategy to monitor the evolution of this problem to adopt health policies to tackle AMR (https://www.who.int/initiatives/glass). Therefore, AMR surveillance systems are paramount in every nation, especially in low- and middle-income countries (LMIC) to better understand and control infectious diseases (4).

Genomic sequencing has allowed a better understanding of AMR (5), and innovative surveillance models has been suggested recently. For instance, metagenomics experiments from sewages have been proposed by international entities to gather AMR information with cheaper and easier standardization than clinical-based AMR monitoring(6).

Given that LMIC contains high rates of infectious diseases (2,3), local and regional AMR monitoring could be established with the sewage approach to assessing AMR genomic regions. Furthermore, Peru represents a country with abundant AMR-associated genes in urban settings (7,8).

Therefore, we present a local genomic assessment of antibiotic resistance genes (ARG) in sewages from two hospitals and urban pipes in a city in Peru, where formal and informal urban settings coexist near rural environments.

## METHODS

### Location and samples

We conducted a metagenomics study in a Peruvian city, namely Huanuco. Our approach used the sewage water collection near two hospitals in 2021: i.e. social security health (ESSALUD) and public non-social security health (Hospital Valdizan) and one from the community, namely urban. These hospitals are the main health service of around 200,000 people from Huanuco. Two samples of one liter (1L) were collected from each sewage on two consecutive days and controlled for environmental factors such as rain or external contamination. After the sample collection, centrifugation, filtering, and storage were conducted.

### Sample, DNA extraction and sequencing

The collection of 250 mL of these sewage waters was collected in sterile tubes for centrifugation and stored at -20 to -80 °C before DNA extraction, using the QIAamp DNA Stool Mini Kit and following the manufacturer’s instructions. DNA Quality and quantification of nucleic acids were performed using the QIAxpert System and gel electrophoresis.

### 16S rRNA gene amplification

To determine the bacterial composition, the V4 variable region of the 16S rRNA bacterial gene was amplified using 16S-515F (GTGCCAGCMGCCGCGGTAA) and 16S-806R (GGACTACHVGGGTWTCTAAT) primers. Libraries were constructed using the dual index approach (9). Sequencing of pooled libraries was carried out using the Illumina MiSeq platform at the University of Minnesota to generate 2x300⍰bp sequences (∼50K reads/sample). 16S rRNA sequences were processed using custom-made Perl scripts and the Qiime2 pipeline. Briefly, raw sequencing data were processed to remove primers and low-quality end reads (Phred quality score⍰<⍰30) using cutadapt. These high-quality reads were considered for denoising, merging, chimera removal, and finally generating unique amplicon sequence variants (ASV) using the Dada2 plugin of Qiime2(10). Representative sequences of each ASV were aligned using MAFFT and phylogenetic trees both rooted and unrooted were constructed using FasTree. Taxonomic assignments of bacterial ASVs were carried out by trained naive Bayes classifiers on reference sequences (clustered at 99% sequence identity) from Greengenes 13_8. For both taxonomic assignments, Qiime2 plugins feature-classifier fit-classifier naïve-Bayes and feature-classifier classifier-sklearn were used. The R statistical software was used to calculate alpha and beta diversity metrics as well as indicator species for treatment groups.

### Shotgun Metagenomics Sequencing

From DNA samples, library prep was conducted using the Nextera XT kit (quarter reaction), and libraries were sequenced on the NovaSeq sequencing platform to produce 2x150 pair-end reads (∼15M reads/sample).

### Bioinformatic pipeline

Filtering and processing of metagenomic sequence data were done using the Kneaddata tool pipeline (https://github.com/biobakery/kneaddata) to remove low-quality reads, primers, and host contamination. The Bowtie2 index of the GRch37 human reference genome was used as the human reference. Trimmomatic parameters within kneaddata were set to 4 base sliding window sizes with only keeping reads with a greater than 20 Phred score. The minimum length of kept samples was also set to at least 90 percent total input read length. Kraken2 was used for taxonomic annotations, using kraken2’s built-in database(11). Taxonomic confidence was set to 0.1. Kraken2’s reports were then processed through Braken for abundance analysis with a minimum length of 100 and at least 10 reads to perform re-estimation(12). Analysis of gene pathway abundance and gene families was performed using Humann3 (http://huttenhower.sph.harvard.edu/humann). Chocophlan and uniref were used as nucleotide and protein databases respectively.

### Antibiotic Resistance Analysis

Analysis was first done using the RGI tool for a gene family, drug class, and resistance mechanism abundance using high-quality, clean sequences(13). DIAMOND was used as the alignment tool for analysis. Secondary analysis of antibiotic resistance was performed using AMRPlus-Plus for annotations of drug resistance mechanisms, resistance type, and functions(14) also leveraging Built-in MEGAres databases.

## RESULTS

The alpha diversity indices (Shannon and Simpson) show that the urban sewage contained more taxa and with higher evenness than the samples from hospitals. The Bray-Curtis index demonstrates the beta diversity of bacterial populations between locations, i.e. hospitals and urban dwellings. Approximately 74% and 79% of the variance between the bacterial taxa and locations was explained in the weighted and unweighted Bray-Curtis index, respectively (Figure 1).

**Figure 1.**
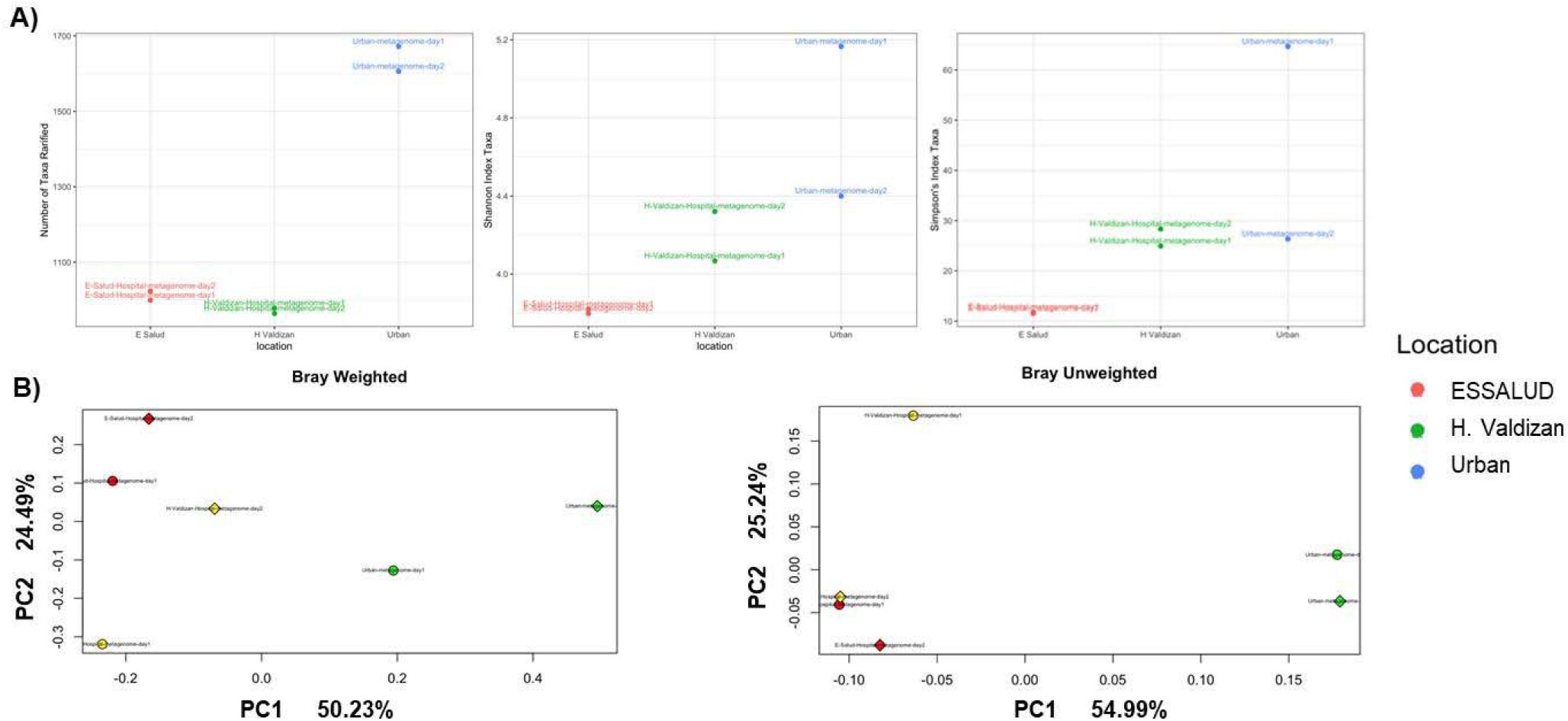
**Alpha and Beta diversity for samples. A) Number of taxa on the left, Shannon (middle) and Simpson’s (on the right) indices. B) Principal coordinate analysis (PCoA) of beta diversity values (unweighted and weighted Bray distances).**

The sewage location influenced the bacterial taxa richness amongst samples. For instance, the ESSALUD hospital shows the most abundant *Prevotella copri* species among all the wastewater samples. Conversely, the urban biospecimen portrayed a higher abundance of *Aeromonas cavie* bacteria, compared to those from hospitals. Even though these proportions were similar between the first and second sample, some differences in taxa abundance were observed between observations for all sites (**Figure 2**).

**Figure 2.**
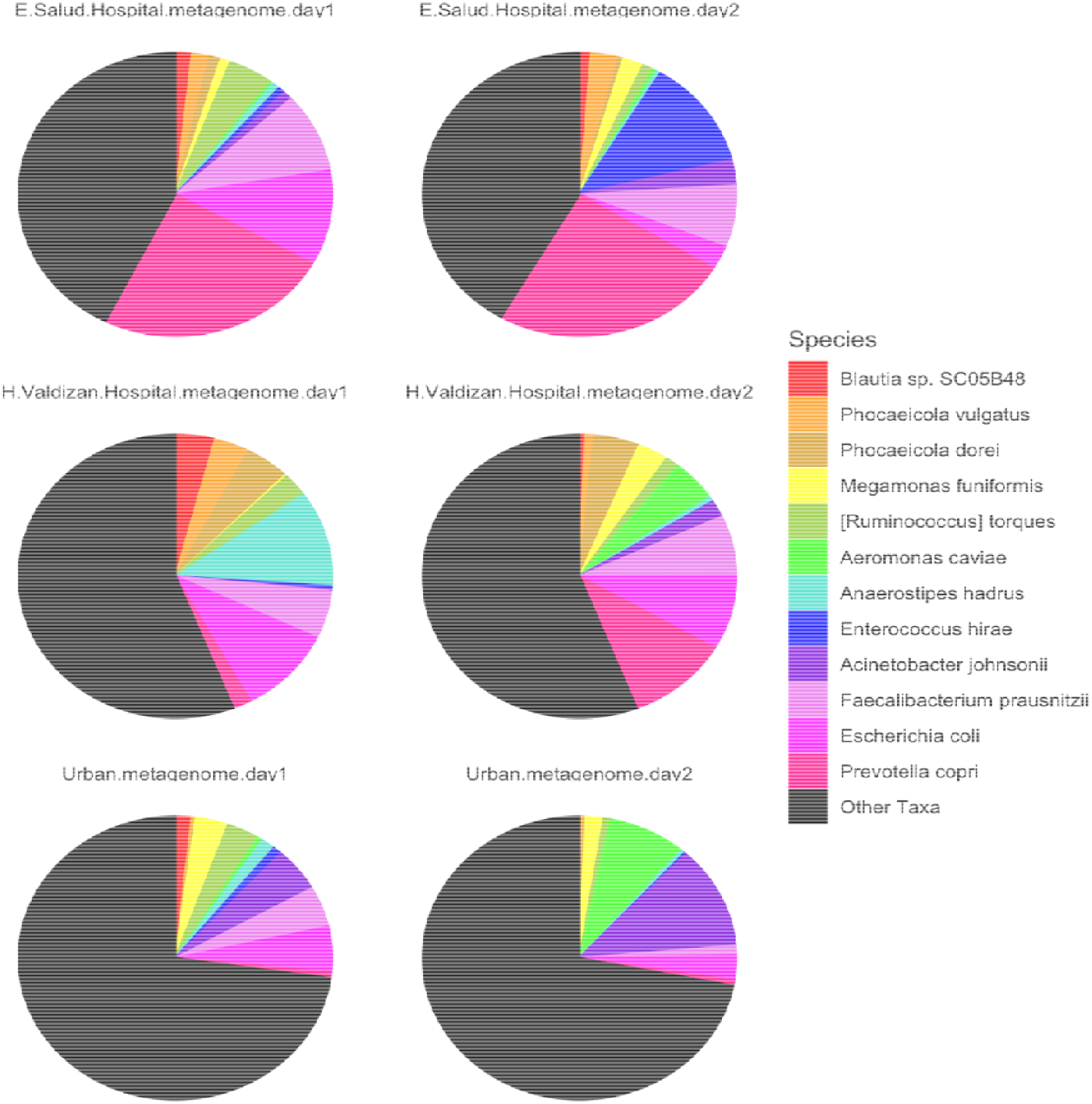
Taxa abundance according to samples and location.

Further analyses demonstrated that the commonest ARG was TEM beta-lactamase or OXA beta-lactamase in all the samples, followed by the major facilitator superfamily (MFS) antibiotic efflux pump, resistance-nodulation-cell division (RND) antibiotic efflux pump, the GES beta-lactamase, the quinolone resistance protein, or the CRX-M betalactamase genes (**Figure 3A**). On the other hand, the samples showed bacteria abundant taxa resistant to Beta-lactam, followed by taxa resistant to cephalosporin, carbapenem, fluoroquinolone and aminoglycoside (**Figure 3B**). The ESSALUD samples showed a similar pattern to urban biospecimens of ARG and resistance to the drug class.

**Figure 3.**
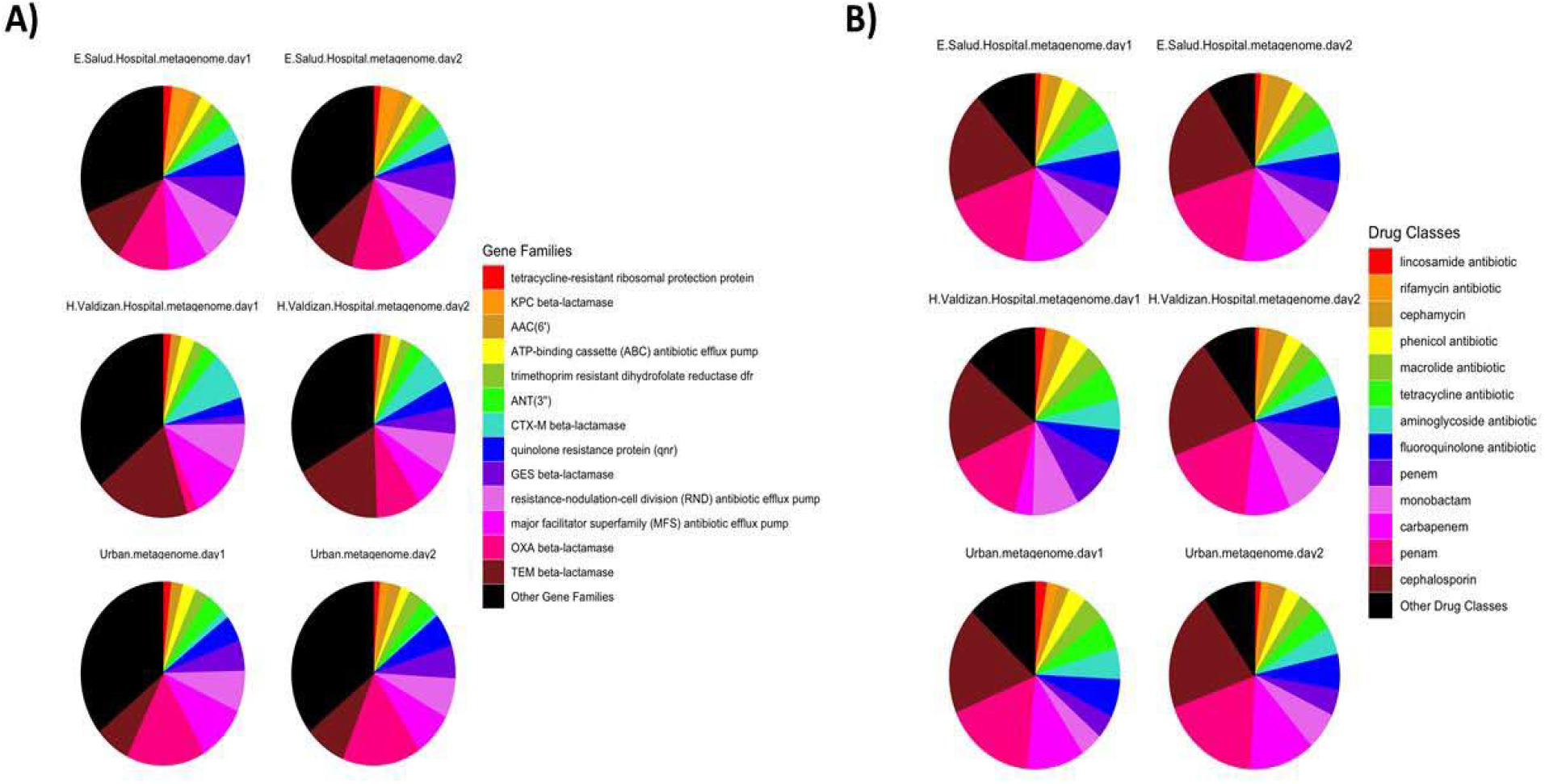
Abundance of Resistance Gene Families (A) and Drug Class Resistance (B)

Figure 4 portrays the resistance according to the drug type, leading the list of an abundance of Macrolides, lincosamides, streptogramins (MLS), followed by bacteria with resistance to aminoglycosides and tetracyclines. It is noteworthy that the sample of ESSALUD contained consistently more bacteria resistant to oxazolidinone than other samples of the other sites. Moreover, the urban sewage depicted more MLS-resistant bacteria than its counterparts. Strikingly, more abundance with rifampin resistance genes was observed in the urban sample than in those near the hospitals.

**Figure 4.**
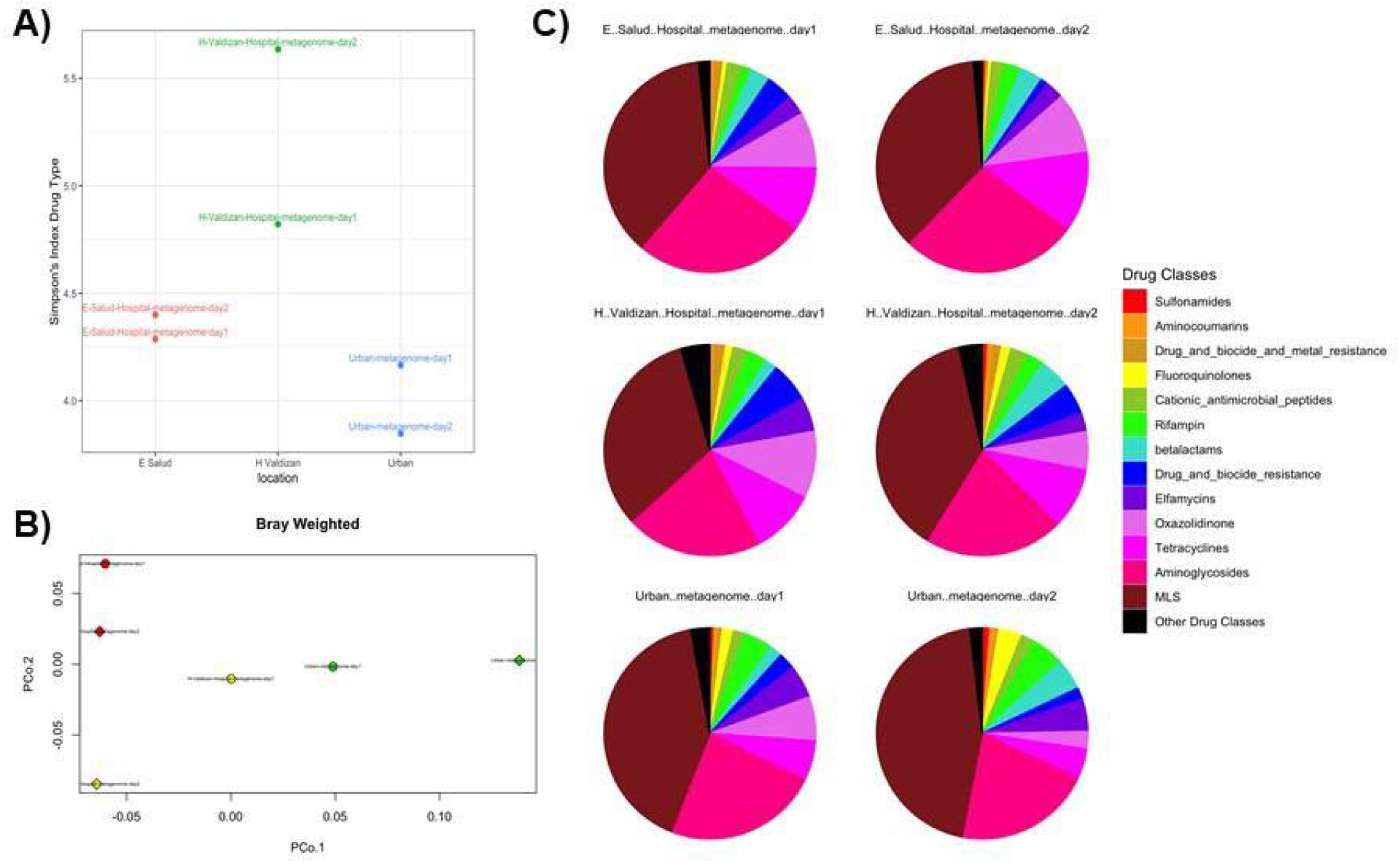
**Antibiotic resistance according to drug type. A) Shannon diversity index to depict alpha diversity (similar to Simpsons’ metric, not shown). B) Beta diversity shown in the Bray-Curtis index and portrayed in the Principal coordinate analysis (PCoA). C) Relative frequencies of bacteria abundance by each type of resistance according to drug type.**

## DISCUSSION

Globally AMR represents a threat that should be monitored in any nation and adapted to specific circumstances. Our study corroborates high levels of resistant genes for beta-lactams, carbapenem, monobactam, macrolides, and even modern antibiotics such as oxazolidinone. Moreover, despite some differences from the wastewater samples near hospitals in this Peruvian city, the major concern refers to the similarity with the urban biospecimen analyzed.

A recent publication, covering 243 cities in 101 countries depicted a localized distribution of ARG according to geographic regions and country, which could further disseminate to other locations. The authors of this report suggested that this could be stable for some periods, but with specific conditions and without control, some bacteria could promote a proliferation across the more extensive areas and promote multidrug bacteria dissemination(8). In this study and its predecessor by o Hendriksen(7), Peru depicted more resistance to macrolides than to beta-lactams, followed by ARG-related to tetracycline resistance. In our study we also found high rates of macrolide-related ARG, but with high rates of ARG for aminoglycosides and beta-lactams.

Antibiotic overuse could be related to the prevalence of infectious diseases, which is common in LMIC(3). Therefore, resistance to macrolides found if our study could be due to the irrational use of azithromycin in the first year of the COVID-19 pandemic (15) or other endemic infectious conditions requiring this sort of treatment (e.g., Helicobacter pylori (16)). Moreover, it is not surprising that we found high levels of resistance to rifampin or fluoroquinolones, as Peru is a country with high rates of tuberculosis (TB) treatment resistance(17).

Results of our project slightly differ from a similar one performed in two sewages of two Hospitals of the Peruvian main city, Lima. In this study, authors suggested that all the bacteria found were multidrug resistant with high rates of ARG associated with extended-spectrum beta-lactamases (ESBL, bla TEM) and carbapenemase genes (ESBL, bla KPC, and bla IMP) (18). Nevertheless, these authors only studied bacteria containing beta-lactamase (bla) related ARG, which could not be fully informative for the resistome of these hospitals in Lima. Conversely, our project used sewage from two hospitals and the urban pipelines to determine the ARG in these three points of the wastewater system in Huanuco, a small city but with geographic, anthropological, and economically diverse relatives sharing a close location.

One of the strengths of our study conceives the concept of ARG assessment in sewages in a local settlement to foster the AMR surveillance system and follow experts’ recommendations (6). Moreover, we evaluated wastewater from effluents of the main healthcare centers in the city and the urban (or community) resistome. This strategy seems to contribute to the assessment of the wastewater system from hospitals and is to our knowledge the first project in a province of Peru following the sewage approach. However, we could not analyze more samples and repeat this process more than once per year. Moreover, we were not able to introduce metagenomics in tap water to assess the resistome directly consumable by humans or animals in this city. It would be crucial to complement our strategy to other methodologies due to a recent study suggesting insufficient effectiveness of Peruvian municipal wastewater treatment plants, where authors quantified moderate concentrations of antibiotics such as clarithromycin, trimethoprim, ciprofloxacin, sulfamethoxazole, and azithromycin (19). Furthermore, as Huanuco is a city with high poverty rates, several parts of the city –if not all of it—resembles an informal urban environment, where several factors such as the crowded houses, farm, and domestic animals, the soil bacteria, amongst others, contribute to AMR (20). This would represent a larger project with a One Health perspective.

## CONCLUSIONS

Metagenomics ARG assessment through the sewage approach seems to be cost-effective in cities such as Huanuco to monitor the antibiotic susceptibility and resistance of bacteria. This could help to establish localized public health policies detecting the community resistome, and assessing the wastewater treatment systems, amongst other activities to foster human, animal, and environmental health in such a biodiverse country as Peru.

## ACKNOWLEDGMENTS

The Fondo Nacional de Desarrollo Científico, Tecnológico y de Innovación Tecnológica (FONDECYT) and BANCO MUNDIAL with contract number 19-2019-BM-INC.INV supported this study. The funders had no role in the study design, the data collection, and interpretation, or the decision to submit the work for publication.

